# Structural and functional analysis of pyocin S8 from *Pseudomonas aeruginosa* : requirement of a glutamate in the H-N-H motif for the DNase activity

**DOI:** 10.1101/2020.04.29.068437

**Authors:** Helena G. Turano, Fernando Gomes, Renato M. Domingos, Maximilia F. S. Degenhardt, Cristiano L. P. Oliveira, Richard C. Garratt, Nilton Lincopan, Luis E. S. Netto

## Abstract

Multi-drug resistance (MDR) is a serious threat to global public health, making the development of new antimicrobials an urgent necessity. Pyocins are protein antibiotics produced by *Pseudomonas aeruginosa* strains to kill closely related cells during intraspecific competition. Here, we report an in depth biochemical, microbicidal and structural characterization of a new S-type pyocin, named S8. Initially, we described the domain organization and secondary structure of S8. Subsequently, we observed that a recombinant S8 composed of the killing subunit in complex with the immunity (Im) protein killed the strain PAO1. Furthermore, mutation of a highly conserved glutamic acid to alanine (Glu100Ala) completely inhibited this antimicrobial activity. Probably the integrity of the H-N-H motif is essential in the killing activity of S8, as Glu100 is a highly conserved component of this structure. Next, we observed that S8 is a metal-dependent endonuclease, as EDTA treatment abolished its ability to cleave supercoiled pUC18 plasmid. Supplementation of apo S8 with Ni^2+^ strongly induced this DNase activity, whereas Mn^2+^ and Mg^2+^ exhibited moderate effects and Zn^2+^ was inhibitory. Additionally, S8 bound Zn^2+^ with a higher affinity than Ni^2+^ and the Glu100Ala mutation decreased the affinity of S8 for these metals as shown by isothermal titration calorimetry (ITC). Finally, we describe the crystal structure of the Glu100Ala pyocin-S8DNase-Im complex at 1.38 Å, which gave us new insights into the endonuclease activity of S8. Our results reinforce the possible use of S8 as an alternative antibiotic for MDR *Pseudomonas aeruginosa* strains, while leaving commensal human microbiota intact.

## Introduction

Increasing rates of antibiotic resistance among Gram-negative pathogens such as *Pseudomonas aeruginosa* represent a serious risk to worldwide health (1, 2). The prevalence of multi-drug-resistant (MDR) strains capable to inactivating a broad array of antibiotics has stimulated the development of alternative therapies (3–5). Bacteriocins produced by *P. aeruginosa*, termed pyocins, represent an emerging antimicrobial option for the treatment of infections caused by MDR strains (6, 7). These molecules are selective protein antibiotics produced by some bacterial species to kill their closely related strains during intraspecific competition (8, 9). Their potency and selectivity towards a narrow-spectrum of *P. aeruginosa* strains make pyocins an attractive antimicrobial alternative to combat MDR strains, while leaving commensal human microbiota intact (10).

Based on their structure, pyocins can be classified into three types called R, F and S. While R/F-type pyocins are high-molecular-weight complexes that resemble phage tails, S-type pyocins are binary protein complexes, comprising a large portion that harbors the killing function and a smaller immunity portion (Figure 1A) (8, 11). The immunity protein protects the pyocin-producing strains by binding with very high affinity (*K*_d_ ≈ 10^−14^ M) and specificity, thereby inactivating the killing protein (12).

**Figure 1.**
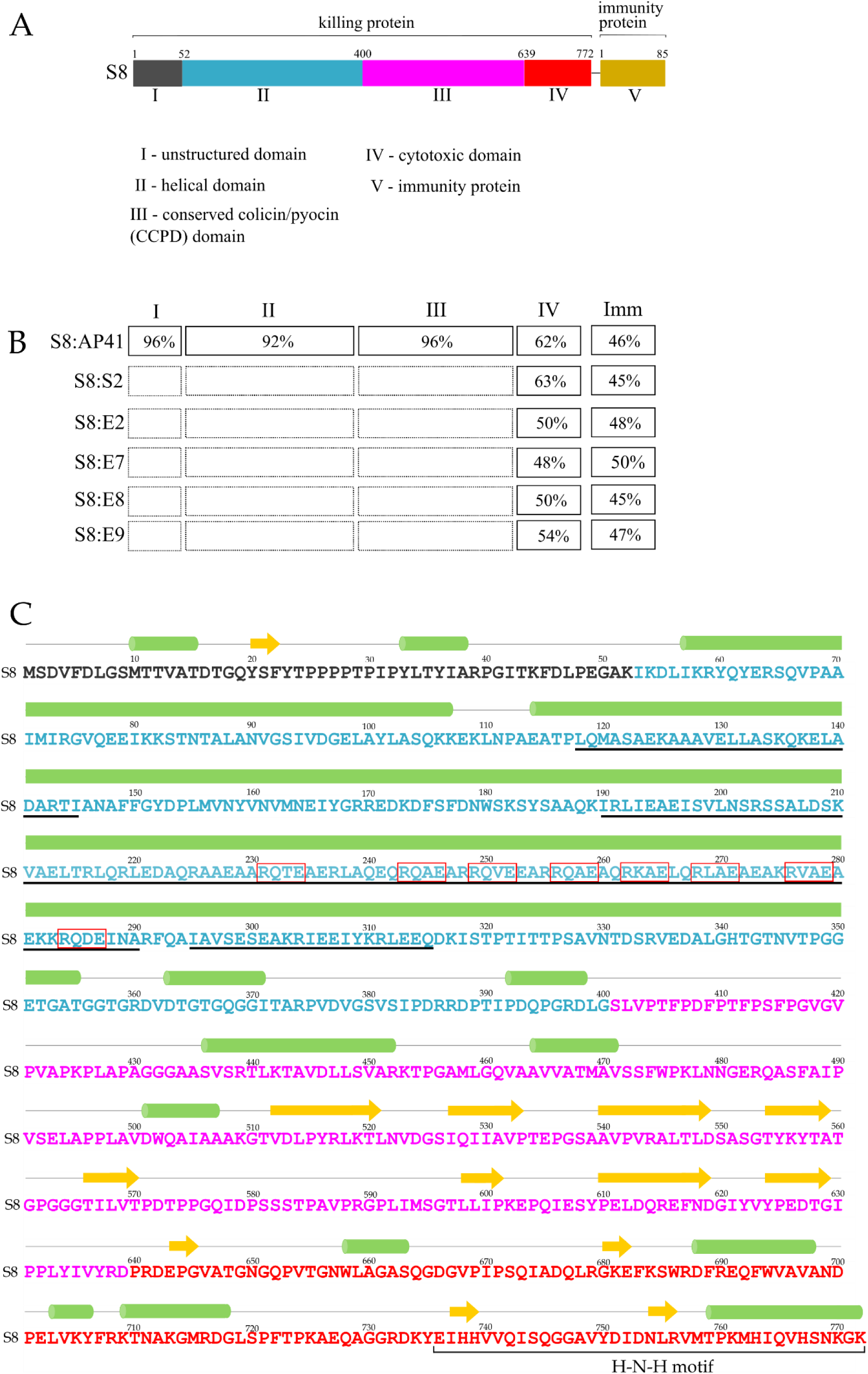
Structural organization of pyocin S8. A - Proposed multi-domain architecture of pyocin S8 based on similarities with others pyocins. The S8 pyocin is a binary protein complex consisting of a large protein that harbors the killing function and a smaller immunity protein. B - Amino acid identities (%) among pyocins and colicins containing the conserved H-N-H-endonuclease motif. C - Predicted secondary structure of pyocin S8 using JPred4 software (45). The amino acid sequences of the different domains are colored according to their respective color as showed in A. Arrows and cylinders represent strand and helices, respectively. The coiled-coil structures are indicated by black underlines. Residues highlighted by red boxes denote the clusters of repeated amino acid sequences R(Q/X)(A/X)E present inside the coiled-coil structure The H-N-H motif are highlighted by a black bracket.

Generally, both killing and immunity proteins are encoded by two separate genes organized in an operon. The transcription of pyocin genes is regulated by the coordinated action of the proteins PrtN, PrtR and RecA (13). PrtN is a transcriptional activator that induces the expression of pyocin genes in response to stressful conditions that causes DNA damage. Under normal conditions, the transcription of PrtN is inhibited by the repressor PrtR. Under stressful conditions, the nucleoprotein filament formed by RecA binding to single-stranded DNA induces de autocleavage of PrtR, leading to the derepression of PrtN and subsequent pyocin production (13–15).

Following their synthesis, the S-type pyocins are released into the extracellular medium as stoichiometric killing/immunity protein complexes. To gain access into their intracellular targets, the S-type pyocins initially parasitize several nutrient uptake pathways that are important for cell viability under nutrient-limited conditions. This process is mediated by the so-called N-terminal receptor-binding domain that binds a specific receptor on the surface of the target cell (16–20). The N-terminal domain then interacts with several components of the proton motive force (PMF)-linked systems (Tol-Pal or Ton systems) in the inner membrane (IM) that span through the periplasm to mediate the pyocin transport across the outer membrane (16, 18, 20). The so-called translocation domain participates in this process but the details still remain poorly understood (19). Some S-type pyocins need also to cross the inner membrane to interact with a cytoplasmic target. However, again, mechanisms underlying this process have been only poorly described. There is evidence that only the C-terminal cytotoxic domain enters the cytoplasm with the immunity protein being released into extracellular medium (21–23). Once inside the cytoplasm, the cytotoxic domain exerts its lethal effect promoting cell killing through a range of mechanisms.

Different S-pyocin cytotoxic domains have been described, most of them displaying nuclease activities towards DNA, rRNA or tRNA, reviewed in (8). DNase domains of pyocins S1, S2 and AP41 bear a conserved H-N-H-endonuclease motif as their catalytic core (24). This H-N-H-motif is also found in colicins, which are well-studied DNases from *Escherichia coli* as well as in other enzymes widely distributed in all kingdoms, serving a variety of functions including homologous recombination, DNA repair, mobile intron homing and apoptosis (25, 26). In the case of pyocins and colicins, the H-N-H motif coordinates a single divalent metal ion and this group catalyzes random cleavages of the bacterial genome (12, 26–29).

The H-N-H motif is composed of 32 amino acids located at the C-terminus of the cytotoxic domain, adopting a topology similar to that of a zinc finger, containing two antiparallel β-strands and one α-helix (ββα-Me) with a single metal ion sandwiched between them (30–32). Several conserved residues of the H-N-H motif play important roles in metal ion binding and DNA hydrolysis. Consequently, substitution of these residues by alanine abolishes the colicin activity *in vivo* (26). Curiously, some of these mutants retain their ability to bind transition metal ions *in vitro* and are able to catalyze DNA cleavage (26). Although the H-N-H motif had been extensively studied in colicins (27), structural information on this motif in pyocins is still scarce.

We have isolated a novel S-type pyocin displaying potent antibacterial activity against MDR *P. aeruginosa* isolates, demonstrating its potential as a protein antibiotic (33). This uncharacterized pyocin, previously designated as S8 by *in silico* analysis (8), has a cytotoxic domain with metal-dependent DNase activity based on an H-N-H motif. Here, we report an in depth characterization of pyocin S8 using multiple biophysical approaches, which include the elucidation of its crystal structure. We also demonstrate the essential role of the highly conserved glutamic acid (Glu100) of the H-N-H motif for pyocin activity.

## Results

### Amino acid sequence identities and domain organization of pyocin S8

We have previously shown that pyocin S8 displays a wide spectrum of killing activity against distinct MDR *P. aeruginosa* strains (33). Pyocin S8 is synthesized as a binary protein complex composed of a large killing subunit that contains 772 amino acids and its cognate smaller immunity protein with 85 amino acids. Similar to other S-type pyocins, the S8 large subunit has a multi-domain organization (Figure 1A). Interestingly, pyocin S8 shares a high amino acid sequence similarity with pyocin AP41, with the exception of the cytotoxic and immunity domains (Figure 1B). In the receptor binding and translocation domains, the amino acid sequence identities are around of 95%, suggesting that these two pyocins are translocated into the target cells by similar mechanisms (Figure 1B).

The C-terminal cytotoxic domain of pyocin S8 bears the H-N-H endonuclease motif that is widely conserved (Figure S1 A) and constitutes the core of their catalytic sites. Although the H-N-H motif is conserved among H-N-H endonucleases, there is still a high level of sequence diversity within the remainder of the cytotoxic domain, especially in the region involved in the interaction with the immunity protein (IPE - immunity protein exosite) (Figure S1 A and B). The cytotoxic domain of pyocin S8 shares about 60% amino acid identity with the cytotoxic domains of pyocins S2 and AP41. The amino acid identities among the cytotoxic domains of pyocin S8 and colicins E2, E7, E8 and E9 are around 50% (Figure 1B).

The first 50 amino acids of pyocin S2 is rich in proline and glycine residues (18, 20), suggesting that this region lacks regular secondary structure. Therefore, this segment is probably unstructured and is followed by a helical domain with a coiled-coil structure. Sequence analysis of pyocin S8 revealed that its N-terminal region is very similar to that of the pyocin S2, containing a proline-rich region (residues 25-32), followed by a coiled-coil structure (Figure 1A and C). Within the helical domain, there is a long cluster of α-helices that contain clusters of amino acid R(Q/X)(A/X)E repeats (Figure 1C, red rectangles). The function of these sequences that were also identified in pyocin AP41 (34) is still unknown.

### Killing spectrum of pyocin S8

The purification of pyocin S8 from *P. aeruginosa* cells is laborious, yielding low amounts of the protein. Therefore, we cloned the genes encoding the entire killing subunit and its cognate immunity protein (ImS8) (Figure 1A) into an *Escherichia coli* expression vector. As ImS8 binds to the killing subunit very tightly, these two subunits were co-purified by nickel affinity chromatography due to the presence of a His6-tag at the C-terminal end of the immunity protein (Figure 2A).

**Figure 2.**
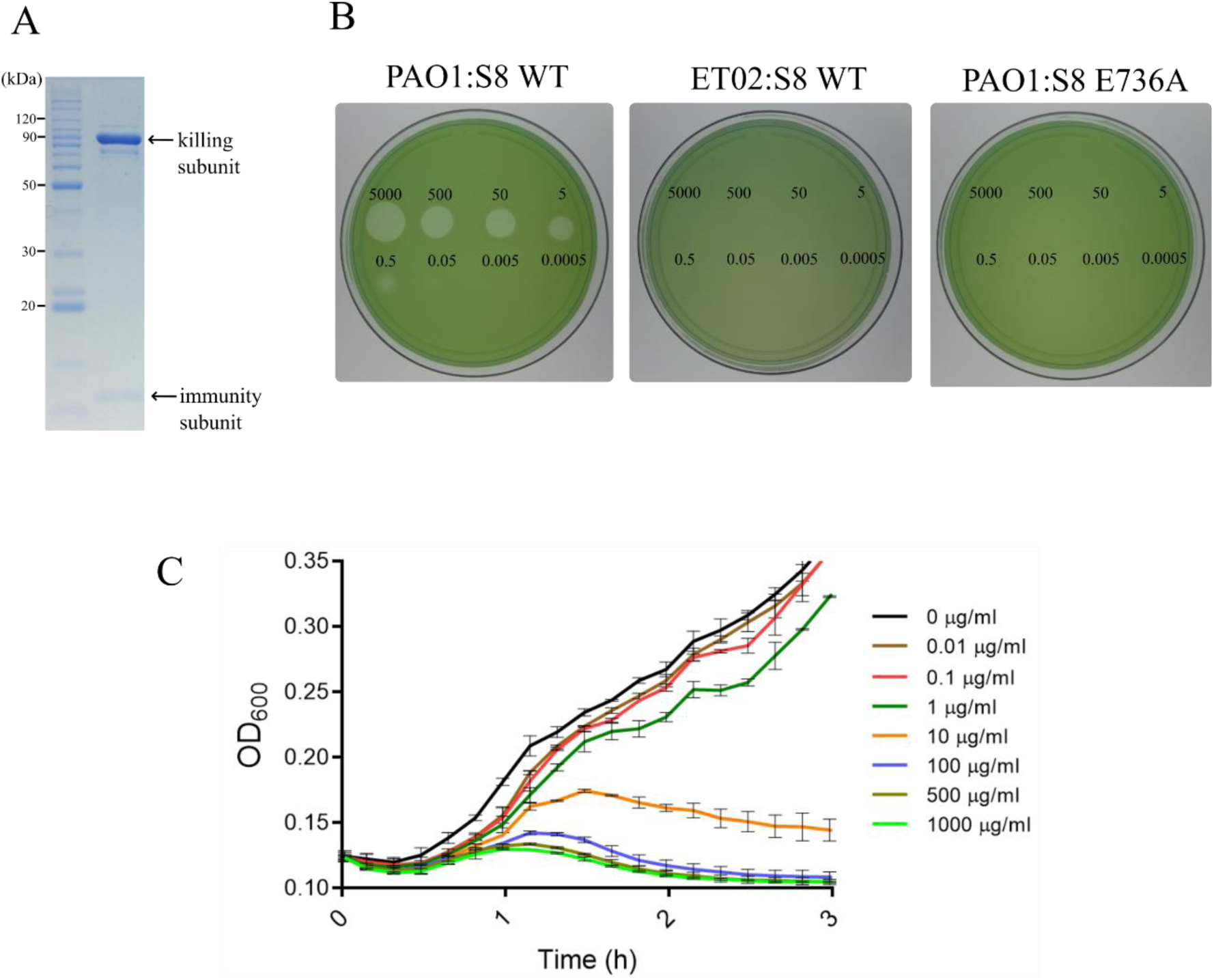
Killing activity of the recombinant pyocin S8. A - SDS-PAGE of purified pyocin S8-ImS8 complex. B - Determination of pyocin S8 activity using an overlay spot plate method. A purified pyocin S8 (starting concentration at 5000 µg/ml) was 10-fold serially diluted and spotted onto the Mueller Hinton agar plates, containing a growing layer of *P. aeruginosa* strains. Clear zones indicate cell death. C - Effect of pyocin S8 on the growth of PAO1 strain. The growth of the cultures in the presence of different concentrations of pyocin S8 was followed at OD_600nm_.

Next, we analyzed the ability of recombinant pyocin S8 to kill *P. aeruginosa* cells. As the PAO1 strain does not produce the pyocin S8 (8), it was employed as a sensitive strain in the killing activity assays. As expected, the S8-ImS8 complex was highly active against the PAO1 strain, as confirmed by clear zones of no growth around the spotting area (Figure 2B, left panel). In contrast, the ET02 strain was completely resistant against pyocin S8, even at the highest concentrations tested (Figure 2B, center panel). This finding was expected, as the ET02 strain contains a gene encoding ImS8 (33), which can bind to the cytotoxic domain and thereby neutralizes its endonuclease activity.

We also observed that the Glu736Ala mutation rendered pyocin S8 inactive against the PAO1 strain (Figure 2B, right panel). These results indicated that Glu736, which is within the DNase active site, is essential for the endonuclease activity.

To measure the pyocin S8 killing activity more precisely, liquid cultures of PAO1 strain were incubated with different amounts of this recombinant protein and its effect on bacterial growth was monitored. The growth of bacterial cultures was not affected at concentrations of pyocin up to 1 µg/ml (Figure 2C). At 10 µg/ml concentration, the growth was severely affected. At concentrations above 100µg/ml, the growth curves displayed an interesting feature: initially, the culture turbidity increased slightly but after 90 min it rapidly decreased to levels lower than observed at the beginning of the curve. These results indicated that pyocin S8 induced cell lysis after a 90 - minute period.

### Endonuclease activity of pyocin S8

In order to test the endonuclease activity of pyocin S8, only the cytotoxic domain of the killing subunit (domain IV in the figure 1A) was utilized in this assay since it is predicted that only this domain penetrates the cytoplasm of the target cells to induce death. The sequence numbering of the cytotoxic domain (referred as S8 DNase domain hereafter) was defined by the alignment of the C-terminal amino acid sequence of pyocin S8 with the DNase domain of the well characterized colicin E9 (30, 35) (Figure S1A). Thus, to facilitate the comparison with other H-N-H endonucleases, we refer hereafter to the sequence positions considering only the E9 DNase domain.

Since the S8 DNase domain cannot be overexpressed without ImS8 due to elevated toxicity for the host strain, both proteins were co-purified as a heterodimeric complex by nickel affinity chromatography employing the C-terminal His6-tag on the immunity protein (Figure 3A, lane 1). Subsequently, the S8DNase domain was separated from ImS8 by treatment with guanidine hydrochloride, followed by another nickel affinity chromatography step to specifically remove ImS8, which remained immobilized in the column (Figure 3, lanes 2 and 3 respectively). Finally, the S8DNase domain was refolded after many cycles of dialysis to remove the denaturant agent (guanidine hydrochloride).

**Figure 3.**
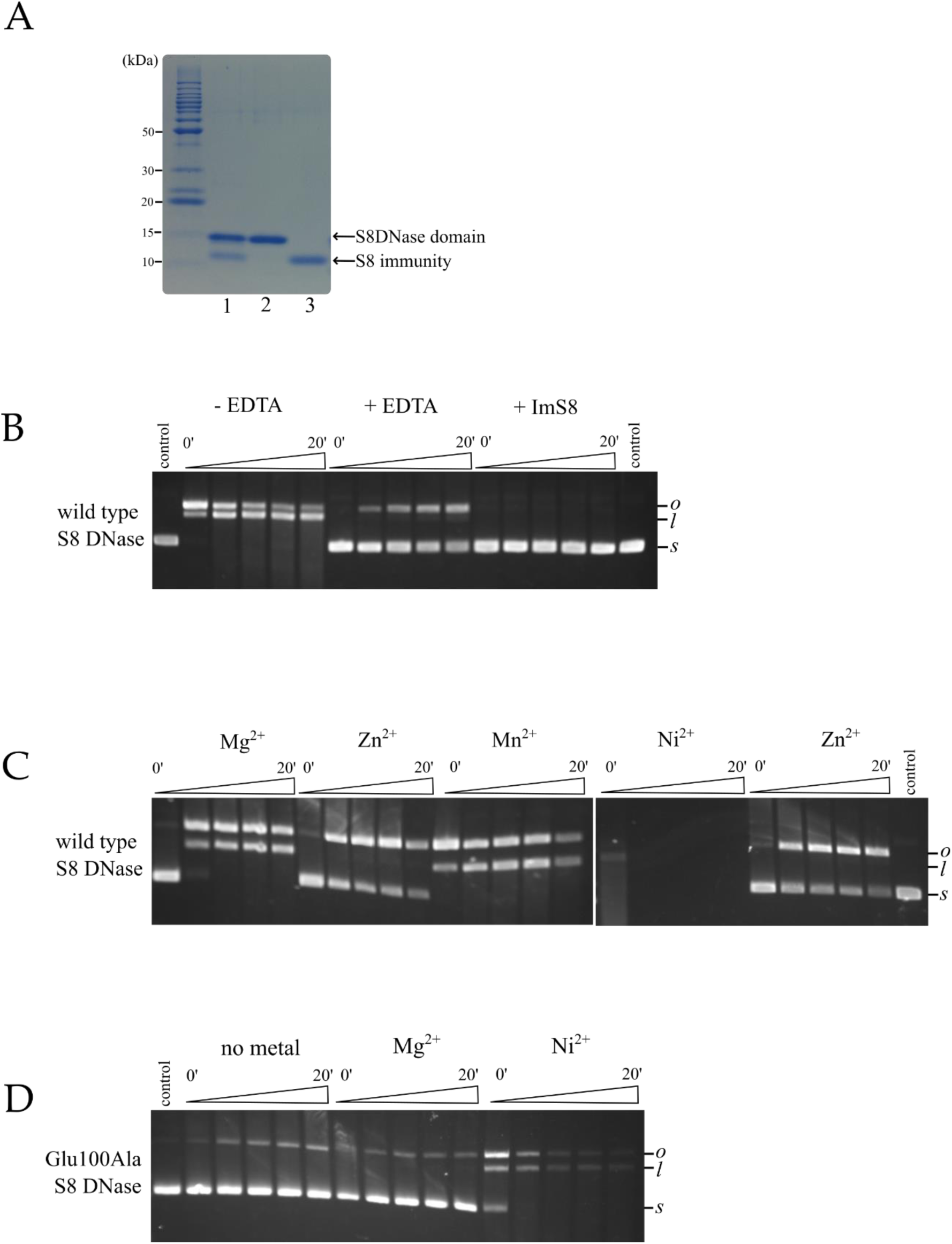
The metal-dependent endonuclease activity of S8 DNase domain. A - SDS- PAGE of purified S8DNase domain and its cognate immunity protein. B - Plasmid nicking assay showing that the endonuclease activity of S8DNase domain is largely reduced by incubation with 20mM EDTA (please observe that the band corresponding to supercoiled DNA (*s*) remained intense even after 20 minutes of reaction). The residual endonuclease activity is completely abolished by the presence of immunity protein. C - recovery of the endonuclease activity of the EDTA-treated wild type S8DNase domain by supplementation with different metal ions at 10mM concentration. D - endonuclease activity of the EDTA-treated Glu100Ala S8DNase as assayed in C. *s*, supercoiled DNA; *o*, open-circle DNA; *l*, linear DNA; control, pUC18 plasmid DNA assayed in the absence of protein.

The ability of the S8 DNase domain to cleave a supercoiled DNA (pUC18 plasmid) was measured. In our experimental conditions, all of the supercoiled (*s*) plasmid was converted into the open (*o*) or linear (*l*) forms (Figure 3B, -EDTA). In contrast, we could observe the band corresponding to the supercoiled form of the plasmid even after 20 minutes of reaction when the S8DNase domain was treated with EDTA, indicating the participation of metal ions in the endonuclease activity (Figure 3B, +EDTA). As expected, the addition of the immunity protein completely abolished the endonuclease activity (Figure 3B, +ImS8).

Next, the metal-specificity of this endonuclease activity was analyzed (Figure 3C). The addition of Mg^2+^ and Mn^2+^ ions reactivated the endonuclease activity of the EDTA-treated S8DNase domain. In contrast, addition of Zn^2+^ did not support the endonuclease activity, even after 20 min of incubation. Indeed, the time course of supercoiled DNA disappearance in the Zn^2+^ supplemented sample was similar to the EDTA-treated S8 DNase domain (Figure 3B, +EDTA). On the other hand, Ni^2+^ induced the highest endonuclease activity, degrading the pUC18 DNA to a much greater extent and precluding the detection of any bands whatsoever. Therefore, the DNase activity is very dependent on the metal ion used in the assay with the highest activity associated with Ni^2+^ and weaker activities associated with Mg^2+^ and Mn^2+^. Similar results have been reported previously for pyocin AP41 (12).

The Glu100Ala S8 DNase mutant (corresponding to Glu736Ala considering the full killing subunit sequence) was not capable to convert supercoiled DNA into linear forms in the presence of Mg^2+^ (Figure 3D), indicating the involvement of the H-N-H motif in this activity. Strikingly, in the presence of Ni^2+^, the mutant exhibited appreciable activity, although considerably lower than the wild type S8DNase. In contrast, the substitution of glutamic acid to alanine in colicin ColE9 from *Escherichia coli* abolishes its endonuclease activity in the presence of nickel (26).

### The affinity of S8 DNase-metal complexes

Since we observed distinct effects of metals on the DNAase activity, we decided to investigate metal-protein interactions by isothermal titration calorimetry (ITC). The stoichiometry of Ni^2+^ and Zn^2+^ binding to S8 DNase domain was 1:1 (Figure 4A) with a substantial enthalpic contribution (>20 kcal/mol), indicating to be a highly favorable event. The dissociation constants for both Zn^2+^ and Ni^2+^ were calculated (Table 1) and indicated that S8 DNase binds Zn^2+^ with an affinity approximately 26-fold higher than Ni^2+^. No binding of Mn^2+^ and Mg^2+^ to the S8 DNase domain was detected (data not show), which is agreement with previous work on the ColE9 DNase domain (36). Possibly, the presence of DNA is required for the S8 DNase domain to bind Mn^2+^ and Mg^2+^ (36).

**Table 1.** Thermodynamic parameters for metals ion binding to the S8 DNase domain determined by ITC experiments. ITC profiles were fitted using a set of sites with same constant affinity model to obtain values for the equilibrium dissociation constant (*K_d_*), ligand binding sites (*n*), and enthalpy change (Δ*H*). Data in parentheses are standard errors from duplicates experiments.

| Experiment | $K_d$ (nM) | $n$ | $\Delta H$(kcal/mol) |
| --- | --- | --- | --- |
| S8 + Zn<sup>2+</sup> | 26.5 (2) | 0.91(2) | -20.8(1) |
| S8 + Ni<sup>2+</sup> | 689 (50) | 0.92(2) | -22.0(6) |
| S8 mutant + Zn<sup>2+</sup> | 508 (20) | 0.96(2) | -17.1(5) |
| S8 mutant + Ni<sup>2+</sup> | 2170 (100) | 0.92(4) | -15.0(9) |

**Figure 4.**
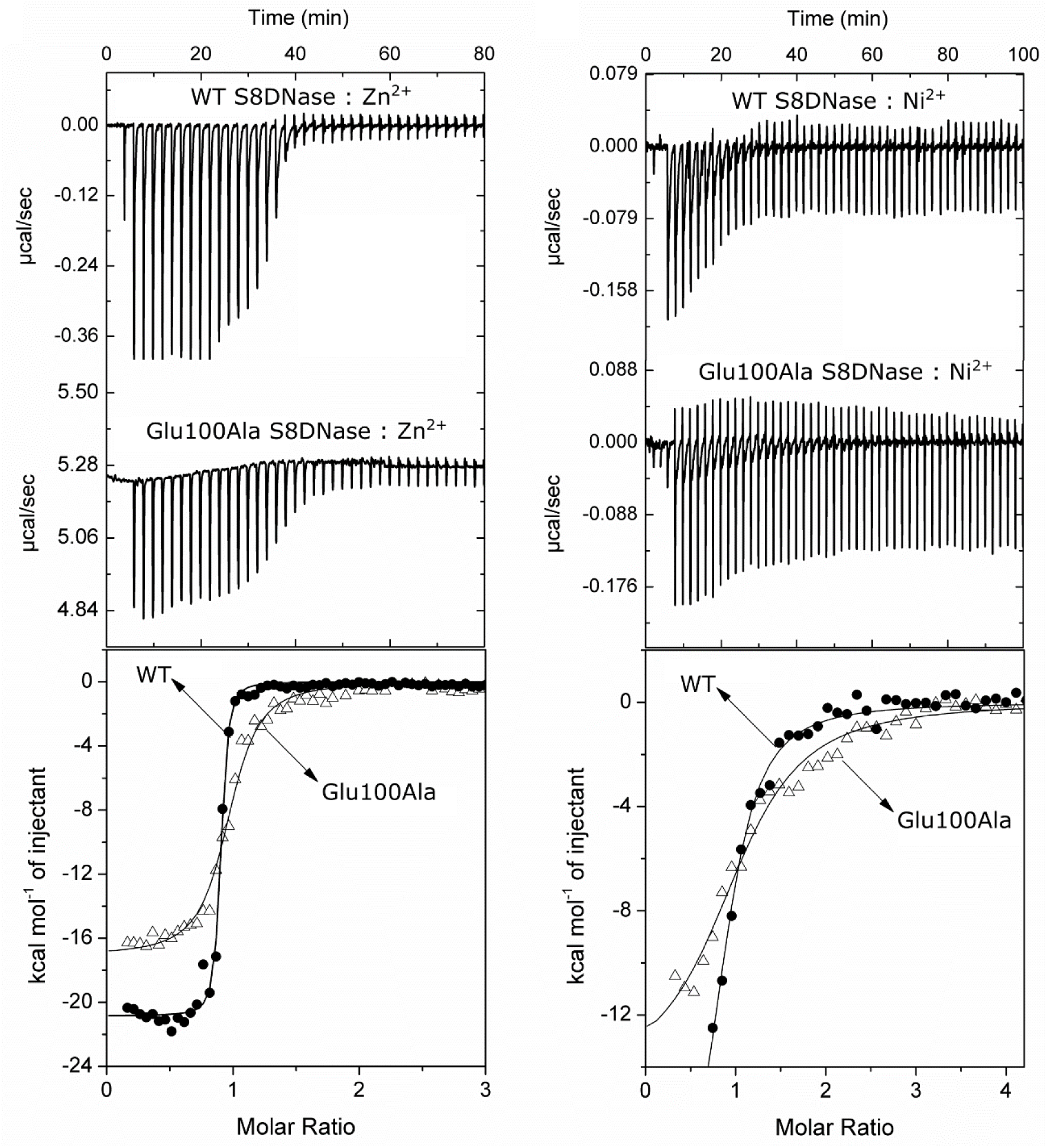
Isothermal titration calorimetry (ITC) for metal binding to S8 DNase domain. *Upper two panels*, ITC thermograms response for the titration of metals (1mM final concentration) into 27 µM of the indicated proteins in 200 mM NaCl, 20 mM Tris-HCl, pH 7.5 buffer at 25°C. Data from two independent experiments were fitted to a simple n-independent binding sites, non-cooperative binding model. *Lower two panels*, heat profile from peak integration of the ITC thermograms of the WT and S8DNase Glu100A mutant, respectively, with the fits to the n-independent binding sites model in black lines. The corresponding thermodynamics parameters are given in Table 1.

The metal-protein affinity of the Glu100Ala S8 DNase mutant was also measured (Figure 4B). The mutant was still able to bind both Zn^2+^ and Ni^2+^ metal ions, even though with a reduced affinity compared with wild-type protein. Possibly, the lower affinity of the S8 Glu100Ala mutant for metals is associated with the fact that this protein could not kill bacterial cells (Figure 2B).

### The crystal structure of the S8 DNase-immunity complex

As described above, the Glu100Ala S8 DNase mutant binds transitions metal ions (Figure 4) and it is able to catalyze DNA hydrolysis in the presence of Ni^2+^ (Figure 3), although with considerably lower efficiency than the wild type protein. Furthermore, this mutant is completely inactive *in vivo* (Figure 2B). In order to better understand the role of this highly conserved residue in the H-N-H motif, the crystal structure of the Glu100Ala S8 DNase domain in complex with ImS8 (hereafter referred to as Glu100Ala S8DNase-Im) was elucidated at 1.38 Å resolution (Table 2).

**Table 2.**
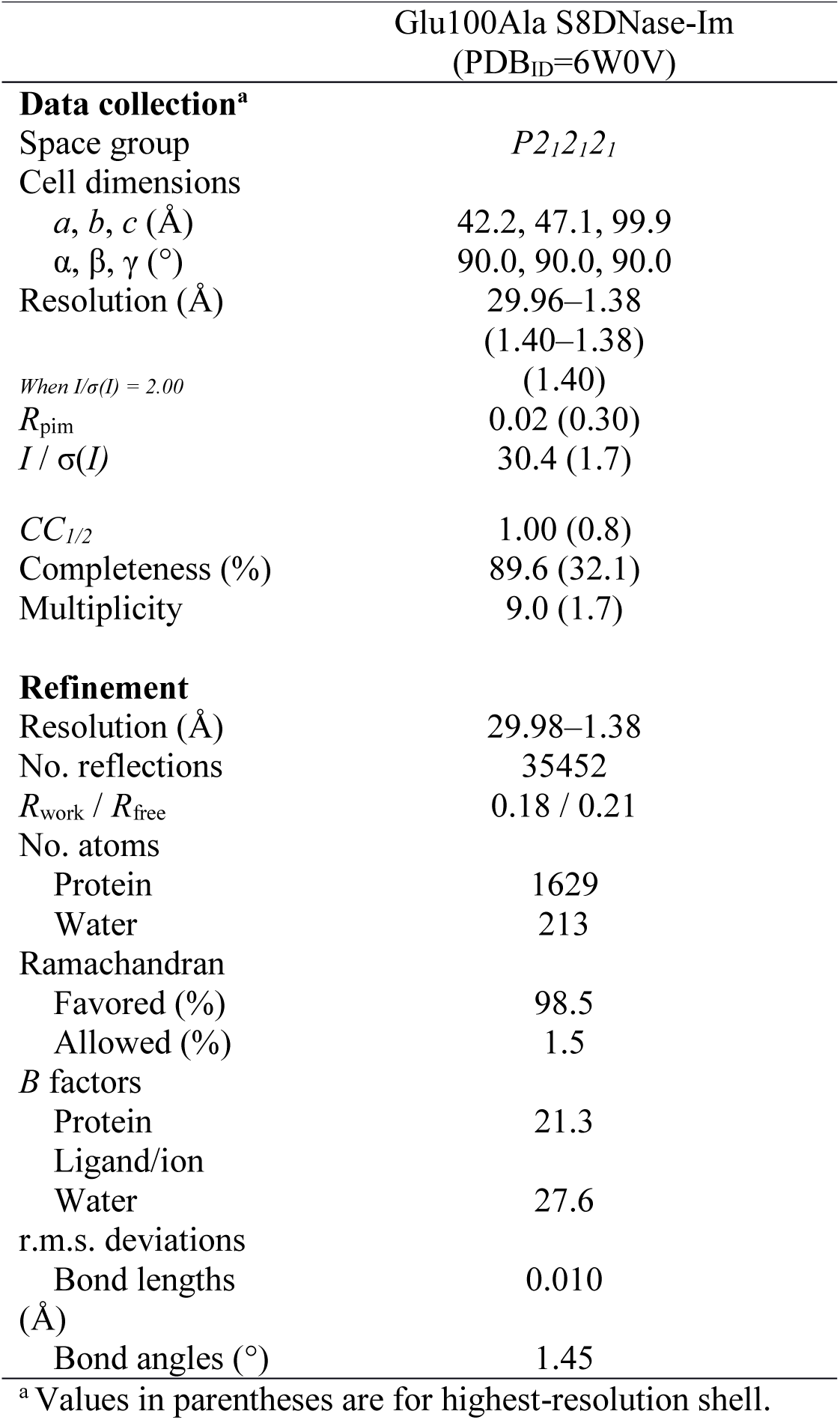
Data collection and refinement statistics.

The refined structure is composed of a 1 cytotoxic: 1 immunity complex in the crystallographic asymmetric unit (Figure 5). The N-terminal methionine and the C-terminal six residues of the DNase domain were not observed in the structure probably because they are unstructured. The overall structure of the Glu100Ala S8DNase is very similar to that of the pyocin AP41DNase (PDB ID: 4UHP) and S2DNase (PDB ID: 4QKO) complexes, with RMSD values for their C^α^ atoms displaying values of 0.72 Å and 0.80 Å, respectively (Figure S2). However, there are notable differences related to the orientation of the amino acids in the active site (see below).

**Figure 5.**
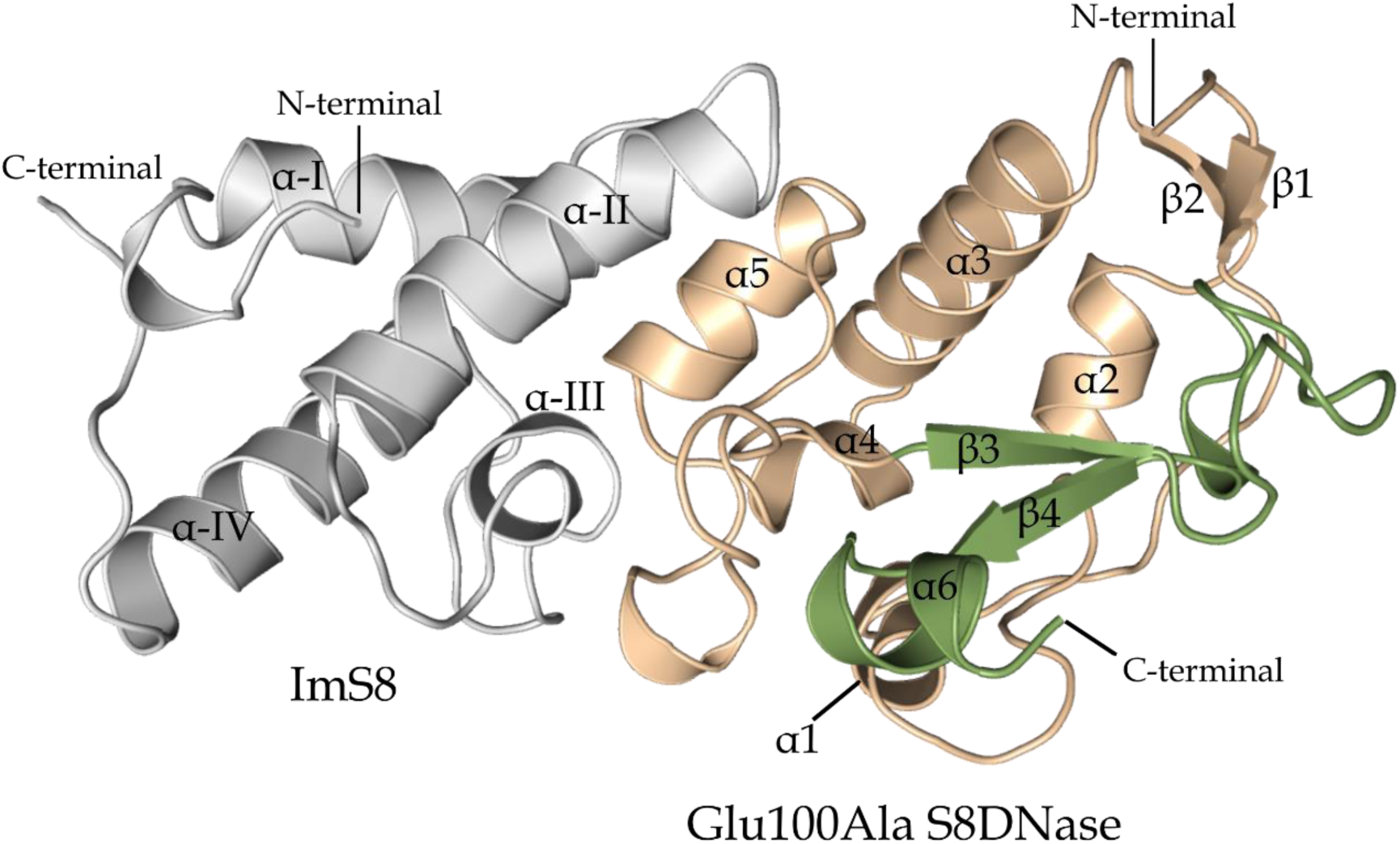
Crystal structure of Glu100Ala S8DNase domain in complex with its immunity protein. The structure of mutant S8DNase in complex with its immunity protein shown in cartoon representation. Glu100Ala S8DNase and its cognate immunity protein are colored light orange/green and gray, respectively. The active site comprising the H-N-H motif are colored green. The S8 DNase domain adopts the typical mixed α/β DNase fold whilst the immunity protein is a four-helix bundle. Figure of protein structure was created using PyMOL.

The structure of the S8 DNase domain is composed of a mixed α/β fold, with its 32 amino-acid H-N-H motif located at the extreme C terminus of the enzyme (Figure 5, region colored green). This motif comprises the active site of the nuclease and adopts a V-shaped architecture, which binds in the minor groove of DNA. ImS8 adopts a four-helix bundle structure and binds to the DNase domain at a region distant from the active site. Although the DNase active site is still exposed in spite of its association with ImS8, somehow this interaction blocks its binding with DNA, probably by steric and electrostatic clashes (12, 30). We detected an inadvertent mutation (Tyr9His) in the α-helix I of ImS8. We believe that this mutation does not interfere with the inhibitory properties of the ImS8, since α-helix I is far away from the cytotoxic–immunity binding interface. Indeed, in the pUC18 cleavage assay, the recombinant immunity protein with the mutation Tyr9His inhibited the ability of S8DNase to cleave DNA (data not shown). Furthermore, the interactions between the DNase and the immunity proteins are also highly conserved among the complexes (12).

### The S8 DNase active site

To further investigate the role of the conserved glutamic acid in the H-N-H motif, we compared the active site of Glu100Ala S8 DNase mutant with others H-N-H DNase structures. Although the amino acid sequences of the structures analyzed are highly similar, they differ considerably in terms of metal ligation. While there are structures showing no metal (apo-form) in the active sites, others are bound to different metals, such as Zn^2+^, Ni^2+^ and Mg^2+^ (12, 28–32, 37, 38) (Figure 6C).

**Figure 6.**
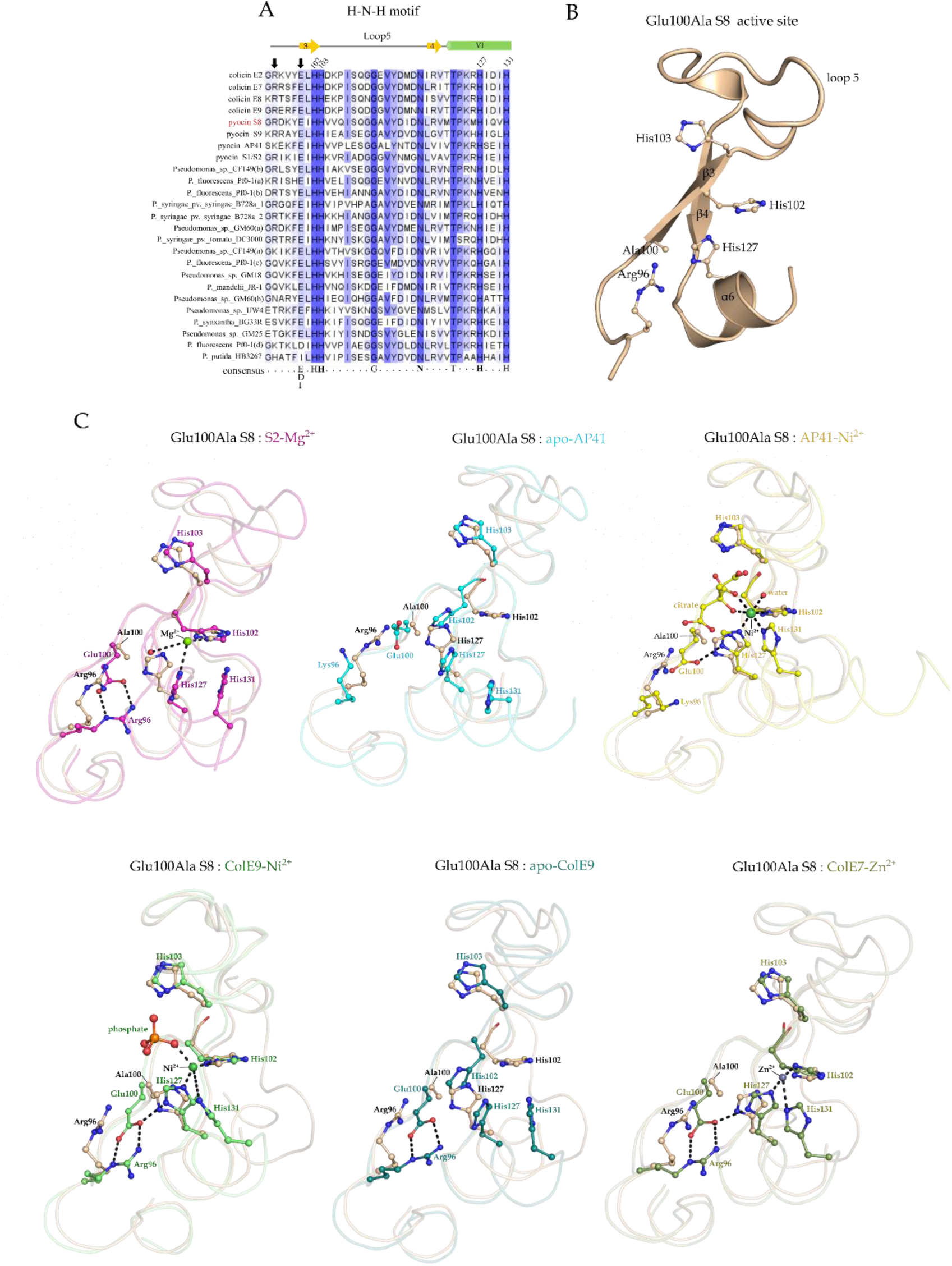
The active site of the H-N-H endonucleases. A - Multiple amino acid sequence alignment, with some residues described in this work highlighted at the bottom as conserved amino acids. The amino acids responsible for the name of the H-N-H motif are highlighted in bold. The numbers at the top refer to the four highly conserved histidine residues. Pyocin S8 name is highlighted in red. The amino acids at positions 96 and 100 (predicted to interact to each other by a salt bridge) are highlighted by black arrows. B - Close view of the Glu100Ala S8DNase active site, shown in cartoon representation with selected side chains shown as sticks. C - superposition of the mutant S8 active site with different H-N-H DNase structures, highlighting key amino acids for DNase activity. The active site is represented in cartoon representation with selected side chains shown as sticks. The structures superposed with Glu100Ala S8DNase were: Mg^2+^-bound S2 DNase-ImS2 structure from 12 (PDB code: 4QKO), apo AP41 DNase-ImAP41 structure from 12 (PDB code: 4UHP), Ni^2+^-bound AP41 DNase structure from 12 (PDB code: 4UHQ), Ni^2+^-bound ColE9 DNase-ImColE9 from 30 (PDB code: 1bxi), apo ColE9 DNase-ImColE9 structure from 39 (PDB code: 1mev) and Zn^2+^-bound ColE7 DNase-ImColE7 structure from 31 (PDB code: 7CEI). Figures of protein structures were created using PyMOL.

The H-N-H motif contains four histidine residues that directly participate in DNA hydrolysis (Figure 6 A, numbered residues). His131 is not observed in the current structure due the absence of electron density for the last six C-terminal residues of the DNase domain (Figure 6 B). His103 is in a very similar conformation to that observed for the corresponding residues in other H-N-H DNase structures (Figure 6C). In fact, this residue plays an important role in catalysis acting as a general base that activates a water molecule for nucleophilic attack of the scissile phosphodiester bond (29, 32). The remaining two histidine residues (His102 and His127) showed slight changes in their conformations in comparison to the other structures (Figure 6C). These histidine residues have been proposed to participate in metal ion coordination and seem to have considerable structural variability depending on the nature of the metal ion present in the structure (29, 32).

The orientation of the sidechains of His102 and His127 are compatible with metal binding being similar to those observed for the Ni^2+^-bound form of AP41 (12) (Figure 6C). However, although weak electron density is visible at the expected metal site in difference maps, it was not possible to attribute this to any particular metal or to establish its occupancy. Therefore, the active site of the S8 DNase domain structure described here is probably predominantly apo (Figure 6B). It is also possible that the weak electron density in the active site could be a PEG molecule (or other component of the crystallization solution), but it was not possible to model it due to steric hindrance effects with the histidine side chains. Thus, although the Glu100A S8 DNase domain is able to bind transition metal ions, the crystal structure presented here appears to be metal-free.

Although Glu100 is not thought to have a direct role in DNA hydrolysis, this residue is highly conserved in the H-N-H motif (Figure 6A, residue highlighted by a black arrow). Indeed, most of the structures show that Glu100 forms a salt bridge with Arg96 (Figure 6C) (12, 29–32, 39). However, there is an exception in the case of pyocin AP41, which contains a lysine in place of arginine. Importantly, in the AP41DNase structures the Glu100 seems to not interact with Lys96 and, consequently, these residues show high mobility (B factor) in the structure (Figure S3).

The Glu100-Arg96 salt bridge has been proposed to play an important role in distorting the DNA substrate, approximating the scissile phosphate towards the metal ion (32). The substitution of Glu100 to alanine, releases constraints on the movement of Arg96, which is consistent with its higher B factor (Figure S3). Therefore, the loss of the Glu100-Arg96 salt bridge probably is the cause of pyocin S8 inactivation.

## Discussion

Here, we describe a biochemical and structural characterization of a new member of the S-type pyocin group. We show that recombinant pyocin S8 is highly effective against the PAO1 strain inducing cell death at concentrations lower than 10µg/ml. This result prompted us to investigate the molecular basis by which this protein shows potent bactericidal activity.

Pyocin S8 is an endonuclease with activity based on the highly conserved H-N-H motif that requires a divalent metal ion as a cofactor. There are some variation regarding the metal ion identity used by H-N-H endonucleases. The most commonly reported metal ion is Mg^2+^, although other divalent cations can also support the H-N-H DNase activity, including Mn^2+^, Zn^2+^ and Ni^2+^ (12, 26, 28, 29, 36, 40, 41). There are conflicting results about the metal requirements for ColE9 and ColE7. While both enzymes are activated by Ni^2+^ and Mg^2+^, Zn^2+^ only activates ColE7, having inhibitory effects on ColE9 (28, 36, 40, 41). In our case, the S8 pyocin exhibited DNase activity when supplemented with Mg^2+^, Mn^2+^ and Ni^2+^, displaying no detectable activity in the presence of Zn^2+^ (Figure 3C). Accordingly, pyocin AP41 DNase displays higher endonuclease activity in the presence of Ni^2+^ or Mg^2+^ ions, showing no detectable activity in the presence of Zn^2+^ (12). Given that a variety of metal ions can support catalysis by H-N-H DNases, the identity of the biologically active metal ion remains a challenge.

Possibly, metals bind H-N-H endonucleases in different ways, activating or inhibiting the histidine residues that are directly involved in the hydrolysis of phosphodiester bonds. The crystal structures of colicin E9 in complex with Zn^2+^-DNA and Mg^2+^-DNA provided some structural insights for the distinct effects of metals on the endonuclease activity of this bacteriocin (32). In the structure of E9 Zn^2+^-DNA, the Zn^2+^ is bound to atoms of the three active site histidine residues (His102, His127 and His131), while in the E9 Mg^2+^-DNA structure, the Mg^2+^ is bound to only two histidine residues (His102 and His127). Therefore, Zn^2+^ binds in the H-N-H motif with high affinity, using His131 as an additional metal-binding site and consequently trapping the enzyme in a catalytically inactive state. Thus, “active” metal ions are those that bind more weakly to the H-N-H motif, leaving His131 in a disengaged state, allowing a water molecule to complete the catalytic cycle (32). In agreement with this model, our ITC analyses showed that Zn^2+^ binds to S8 with higher affinity than Ni^2+^. However, the situation is complicated by the fact that the AP41 Ni^2+^-bound complex shows the metal bound to all three histidine side-chains. Therefore, crystal structures of pyocin S8 in complex with DNA will be required to understand the mechanisms by which the endonuclease activity of this bacteriocin is modulated by metals.

Besides metals, this work provided new insights on the function of the H-N-H motif. Our structure of the S8 DNase–Im8 complex is only the third cognate pyocin DNase–Im protein complex solved and this is the first displaying a mutation of the conserved Glu (Glu100) in the H–N–H motif. As this conserved Glu (Glu100) is essential to the killing activity (Figure 2), the comparison of our structure with the others available provided valuable information. The superposition of H-N-H DNase-Im structures showed that the Glu100-Arg96 salt bridge is highly conserved (Figure 6C). Noteworthy, Arg96 is positioned in a distinct rotamer in our DNase (Glu100Ala) structure, suggesting that Arg96 presents high degree of freedom to assume distinct conformations.

The H-N-H motif constitutes the DNase active site of pyocins and closely related colicins. The current model of DNA hydrolysis by H-N-H endonucleases suggests that the double helix of DNA needs to undergo some distortion to approximate the scissile phosphate close to the metal center of the H-N-H motif (26, 29, 32). This is achieved by positioning the H-N-H motif into the minor groove of DNA, mainly due the insertion of the Arg-96-Glu-100 salt bridge into the groove itself. As a result of these DNA-enzyme interactions, the scissile phosphate is positioned toward the metal ion embedded within the H-N-H motif (32).

In the Glu100Ala mutant, the lack of the salt bridge with Arg96 frees up the latter’s side-chain to the extent that it presents much weaker density than the surrounding residues and appears to have moved away from its canonical orientation. Clearly this ultimately has a knock-on effect which influences the ability of the V-shaped H-N-H motif to interact with DNA and catalyze DNA hydrolysis. The most likely explanation is that the normal role of Glu100 is to correctly orientate the guanidinium group of Arg96 by accepting hydrogen bonds involving N_ε_H and one of the NH_2_ groups. This leaves the remaining hydrogen atoms free to interact with hydrogen-bond acceptors from both the DNA backbone and one of the bases (32). In the absence of such interactions in the Glu100Ala mutant, the distortion of DNA probably does not occur, which could explain the loss of the killing activity. However, the determination of the crystal structure of S8 pyocin in complex with DNA substrate is required to confirm this idea.

As described before, the H-N-H motif is composed of four conserved histidine residues that play important roles in DNA hydrolysis. The role of His131 is still elusive and appears to be involved in the activation of a water molecule to protonate the 3’-oxygen-leaving group of a phosphodiester bond (29). In some cases, this residue can also coordinate metals, as showed in the Zn^2+^-bound DNases structures (28, 31, 32). His102 and His127 residues appear always engaged in metal ion coordination, regardless of which metal is bound to the enzyme. Finally, the His103 activates a water molecule for the nucleophilic attack of the scissile phosphate bond (29, 32). With the exception of the His131, which is disordered in our structure, the other three histidine residues showed similar orientations to equivalent residues in the other DNase–Im structures. Thus, although the putative metal binding sites (represented by the active site histidine residues) were only moderately affected in our structure, no metal ion was found in the active site. Joshi e coworkers (12) have reported the crystal structures for the wild-type pyocin AP41 DNase–Im and pyocin S2 DNase–ImS complexes. The former is in the apo-form whilst the latter, curiously, has been modeled including a Mg^2+^ ion at the active site despite the absence of DNA in this structure. The diversity of the active site configurations observed in the current structures of the H-N-H endonucleases suggest that these enzymes have a dynamic catalytic center, able to adapt to different crystallization conditions.

Bacteriocins such as pyocins and colicins play important roles in shaping bacterial communities and also affecting host-pathogen interactions (15, 42–44). Understanding the molecular mechanisms by which pyocins kill target cells is a promising approach to development new therapeutic approaches for MDR infections caused by *P. aeruginosa*. Moreover, while these proteins could be successfully used in the treatment of infections caused by specific *P. aeruginosa* MDR strains, a minimal collateral damage would be expected to occur to beneficial microbiota species.

## Materials and Methods

### Secondary structure prediction and sequence alignments

The secondary structure elements of pyocin S8 were predicted by using the JPred4 server available in the Jalview program (45). Pyocin protein sequence alignments were performed using Jalview program.

### Plasmid constructions

The genes encoding either the full-size pyocin S8 and its immunity protein were PCR-amplified from ET02 genomic DNA using primers 5’-GGAATTCCATATGAGCGACGTTTTTGACCTTGGA-3’ and 5’-ATGGATCCTTGCCAGCCTTGAAGCCAGGGAG-3’. The PCR product was digested with the NdeI and BamHI and ligated into the corresponding sites of the *E. coli* expression vector pET29b to give pET29b-PyoS8-ImS8, which encodes pyocin S8 operon with a C-terminal His6 tag fused to the immunity protein. To create the substitution of the conserved glutamic acid 736 (glutamic acid at position 100 when considering only the cytotoxic domain) in the H-N-H motif by alanine, pET29b-PyoS8-ImS8 was used as a template for the site-directed mutagenesis (QuikChange XL site-directed mutagenesis kit; Agilent Technologies) to yield pET29b-PyoS8Glu736Ala-ImS8.

The coding sequence for pyocin S8 cytotoxic domain starting in aspartic acid residue at position 6 (residues 642-772 in terms of the full-length pyocin S8 killing subunit) and its cognate immunity protein were PCR-amplified from ET02 genomic DNA using primers 5’-ATTATATCATATGGATGAGCCGGGTGTTGCTACC-3’ and 5’-ATGGATCCTTGCCAGCCTTGAAGCCAGGGAG-3’. The PCR product was digested with the NdeI and BamHI and ligated into the corresponding sites of the pET29b to give pET29b-PyoS8DNase-ImS8. The coding sequence for pyocin S8 cytotoxic domain containing the mutation of glutamic acid at position 100 by alanine and immunity protein were PCR-amplified from pET29b-PyoS8Glu736Ala-ImS8 using primers 5’-ATTATATCATATGGATGAGCCGGGTGTTGCTACC-3’ and 5’-AAGGTACCGCCAGCCTTGAAGCCAGGGAG-3’. The PCR product was digested with the NdeI and KpnI and ligated into the corresponding sites of the pET29b to give pET29b-PyoS8DNaseE100A-ImS8.

### Protein overexpression and purification of pyocin S8

The expression vector encoding the full-size pyocin S8 and its immunity protein was transformed into *E. coli* BL21(*DE3*) pLysS competent cells. Overnight cultures were diluted in 0,5 L of LB broth to an *OD*_600_ = 0.2 and cells were grown at 37 °C in a shaking incubator to an *OD*_600_ = 0.6-0.8. The culture was then shifted from 37°C to 30°C and protein expression was induced by addition of 0.5 mM isopropyl β-D-1-thiogalactopyranoside (IPTG) during a period of 6 h. Cells were then harvested and resuspended in a 20 ml start buffer (20 mM sodium phosphate, 500 mM NaCl, pH 8.0) containing 0.05 mg/ml lysozyme. After incubation at 4°C for 30 min, 500 microliters of a 10x Complete EDTA-free protease inhibitor cocktail (Sigma Aldrich) solution was added to the cell suspension. Cells were lysed by sonication and the cell extract was kept on ice for 20 min during a 1% streptomycin sulfate treatment. Cell debris was removed by centrifugation at 15,000 × *g* at 4°C for 40 minutes, and supernatant was further clarified by filtration using a 0.45 µm pore membrane. The cell-free lysate was applied into a 5 ml Hi Trap Chelating HP column (GE Healthcare) equilibrated in start buffer. After loading, the column was connected into an AKTA TM FPLC system (Amersham Biosciences, GE Healthcare), and washed with 50 ml start buffer containing 50 mM imidazole to remove unbound protein. The bound protein was then eluted by an imidazole gradient (20-500mM). Then, the purified proteins were dialyzed against TN_200_ buffer (20 mM Tris-HCl pH 7.5, 200 mM NaCl) using a HiTrap Dessalting column (GE Healthcare), and the protein was further purified by gel filtration chromatography on a HiLoad 16/600 Superdex 75 or HiLoad 16/600 Superdex 200 column (GE Healthcare) equilibrated in the same buffer. The purity of the proteins were asses by SDS-PAGE and the fractions were pooled pure and concentrated using an Amicon Ultra concentrator (Merck Millipore) with a 10 KDa cut-off. The proteins were divided into small aliquots and stored at −80°C. Prior to use protein concentrations were determined by absorbance at 280 nm using the molar extinction coefficient (ε_280_ = 77240 M^−1^ cm^−1^) obtained using the ProtParam tool.

### Purification of DNase domain from ImS8

Initially the PyoS8DNase-ImS8 complex was overexpressed and purified essentially as described above. Then, the PyoS8 DNase domain was isolated from the PyoS8DNase-ImS8 complex using 6 M guanidine hydrochloride as described in (12). Briefly, the PyoS8 DNase-ImS8 complex at 5.5mg/ml was diluted 10-fold in a denaturing buffer containing 500 mM NaCl, 20 mM Tris-HCl (pH8.0) and 6 M guanidine hydrochloride. The sample was then incubated for 1 h at room temperature with occasional agitation. Pyo S8 DNase domain was separated from its cognate immunity protein by loading the sample onto a 5 ml Hi Trap Chelating HP column (GE Healthcare) equilibrated with denaturing buffer. In this step, the immunity protein was immobilized in nickel affinity resin due to a C-terminal His6-tag. The DNase domain was then concentrated and incubated with 20 mM ethylenediaminetetraacetic acid (EDTA) for 1 h at 25°C to remove adventitious metal ions. The chelating agent and the denaturant agents were removed by extensive dialysis against 50 mM Tris (pH 7.5) buffer. Finally, after concentration and quantification, protein was divided into small aliquots and stored at −80°C. The PyoS8 DNase E100A domain was purified in the same way described above.

### Pyocin sensitivity assays

Activity of recombinant pyocin S8 was assessed by the overlay spot plate method as described previously (12) with a slight modification. Four hundred microliters of a sensitive strain culture at 0.5 McFarland turbidity (1.5 × 108 CFU/ml of bacteria) was added to 6 ml of 0.7% soft agar and poured over onto Muller Hinton or G medium agar plates. Purified pyocin S8 at initial concentration of 5mg/ml was 8-fold serially diluted and three microliters of each dilution was spotted onto the plates and incubated for 16-18 hours at 37°C.

Alternatively, bacterial cultures at *OD*_600_ = 0.1 were incubated with different concentrations of pyocin S8 and grown at 37°C with continuous shaking in a 96 well plate. The effect of pyocin S8 on the growth was monitored by measuring the turbidity of the cultures every 10 min for 3 h using the automated Sinergy™ H1 absorbance microplate reader (BioTek).

### Endonuclease activity assay

The endonuclease activity was evaluated by plasmid nicking assay as described by (12) with some modifications. A supercoiled pUC18 (1 µg) was used as the substrate for the plasmid nicking assay. Assays were performed at 25°C in a buffer containing 50 mM Tris-HCl (pH 7.5) in a final volume of 50 µl. Reactions were started by the addition of PyoS8 DNase domain (previously treated with EDTA) to a final concentration of 0,4 µM. At different time points, aliquots of 10 µL were removed and reaction was stopped by the addition of sample buffer containing 10 mM EDTA. Metal dependence was tested by supplementing different metal ions to a final concentration of 10 mM. The plasmid nicking results were analyzed on 1% agarose gels.

### Isothermal titration calorimetry (ITC)

The ITC experiments were carried out at 25 °C in 20mM Tris-HCl, 200 mM NaCl pH 7.5 buffer using a MicroCal ITC200 (GE Healthcare). The protein solutions (27 µM) were gently loaded into the sample cell consisting of 200 µl working volume. The titration involved 0.3-0.5 µl injections of ligands present in a syringe filled with 39 µl of metal ions solutions diluted in ultra-pure water to a final concentration of 1mM. The measurements were performed by 77 injections of 0.5 µl or 0.3 µl for Ni^2+^ and Zn^2+^, respectively, separated by delay of 120 seconds between each injection. Control dilutions experiments involving injections of ligands into buffer were performed and subtracted from the integrated data before curve fitting. Data were fit to a model with n-sites with same biding affinities using MicroCalTM software (OringTM package) to determine the dissociation constant, enthalpy of binding (*ΔH*), and stoichiometry. These parameters are reported as the mean value obtained from two independent experiments performed on distinct protein productions and the corresponding sample standard deviation.

### Crystalization and data collection

Crystals were grown at 18°C using the hanging-drop vapor diffusion method. The mutant Glu100Ala S8DNase-Tyr9His ImS8 crystals were grown by adding 10mg/ml of 6xHis-protein (diluted in 5mM Tris-HCl pH 7.4 in an equal volume of the reservoir solution containing Sodium Acetate 0.2M, PEG 4000 30% (w/v), Tris-HCl 0.1M pH 8.0. The crystals were all flash frozen in liquid N2 and collected at beamline MX2 at the Brazilian Synchrotron Light Laboratory (LNLS).

### Structure determination and refinement

All data were indexed and integrated with XDS (46) and scaled using AIMLESS in the CCP4 program suite (47). Initial phases were obtained by molecular replacement with PHASER (48) using *Pa*Pyocin AP41 (PDBID 4UHP) as the search model. One copy of a monomer of the 1cytotoxic: 1 immunity complex was found in the asymmetric unit. Models were built in COOT (49) and refined using REFMAC5 (50). Details of the refinement statistics are presented in Table 2.

## Acknowledgments

We thank Simone V. Alves and Thiago G. P. Alegria for their technical support.

## Author contributions

H.G.T and F.G. conceptualization; H.G.T., F.G., R.M.D., M.F.S.D., L.E.S.N validation; H.G.T., F.G., R.M.D., M.F.S.D. investigation; H.G.T., F.G., R.M.D., M.F.S.D., C.L.P.O., R.C.G., N.L. and L. E. S. N. methodology; H.G.T., F.G., L.E.S.N writing-original draft; H.G.T., F.G., R.C.G., L.E.S.N writing-review and editing; H.G.T., F.G., R.M.D., M.F.S.D., R.C.G., L. E. S. N. formal analysis; H.G.T., F.G., R.M.D., M.F.S.D., L. E. S. N. data curation; N.L. resources; C.L.P.O., R.C.G., and L. E. S. N. supervision; L. E. S. N. funding acquisition; L. E. S. N. project administration.

## Funding and additional information

This work was supported by São Paulo Research Foundation (FAPESP) Grant 013/07937-8, Conselho Nacional de Desenvolvimento Científico Tecnológico (CNPq) Grant 573530/2008-4, and Pro-Reitoria de Pesquisa da Universidade de São Paulo (PRPUSP) Grant 2011.1.9352.1.8. HGT, FG and LESN are members of NAP Redoxoma (PRPUSP), and the CEPID Redoxoma (FAPESP).

## References

1. Horcajada JP, Montero M, Oliver A, Sorlí L, Luque S, Gómez-Zorrilla S, Benito N, Grau S. 2019. Epidemiology and treatment of multidrug-resistant and extensively drug-resistant Pseudomonas aeruginosa infections. Clin Microbiol Rev 32:1–52.

2. Shortridge D, Gales AC, Streit JM, Huband MD, Tsakris A, Jones RN. 2019. Geographic and Temporal Patterns of Antimicrobial Resistance in Pseudomonas aeruginosa Over 20 Years From the SENTRY Antimicrobial Surveillance Program, 1997–2016. Open forum Infect Dis 6:S63–S68.

3. Ghosh C, Sarkar P, Issa R, Haldar J. 2019. Alternatives to Conventional Antibiotics in the Era of Antimicrobial Resistance. Trends Microbiol 27:323–338.

4. Heselpoth RD, Euler CW, Schuch R, Fischetti VA. 2019. Lysocins: Bioengineered Antimicrobials That Deliver Lysins across the Outer Membrane of Gram-Negative Bacteria. Antimicrob Agents Chemother 63:1–14.

5. Theuretzbacher U, Outterson K, Engel A, Karlén A. 2019. The global preclinical antibacterial pipeline. Nat Rev Microbiol.

6. McCaughey LC, Ritchie ND, Douce GR, Evans TJ, Walker D. 2016. Efficacy of species-specific protein antibiotics in a murine model of acute Pseudomonas aeruginosa lung infection. Sci Rep 6:30201.

7. Redero M, López-Causapé C, Aznar J, Oliver A, Blázquez J, Prieto AI. 2018. Susceptibility to R-pyocins of Pseudomonas aeruginosa clinical isolates from cystic fibrosis patients. J Antimicrob Chemother 73:2770–2776.

8. Ghequire MGK, De Mot R. 2014. Ribosomally encoded antibacterial proteins and peptides from Pseudomonas. FEMS Microbiol Rev 38:523–68.

9. Michel-Briand Y, Baysse C. 2002. The pyocins of Pseudomonas aeruginosa. Biochimie 84:499–510.

10. Ghequire MGK, De Mot R. 2018. Turning Over a New Leaf: Bacteriocins Going Green. Trends Microbiol 26:1–2.

11. Scholl D. 2017. Phage Tail-Like Bacteriocins. Annu Rev Virol 4:453–467.

12. Joshi A, Grinter R, Josts I, Chen S, Wojdyla JA, Lowe ED, Kaminska R, Sharp C, McCaughey L, Roszak AW, Cogdell RJ, Byron O, Walker D, Kleanthous C. 2015. Structures of the Ultra-High-Affinity Protein-Protein Complexes of Pyocins S2 and AP41 and Their Cognate Immunity Proteins from Pseudomonas aeruginosa. J Mol Biol 427:2852–66.

13. Matsui H, Sano Y, Ishihara H, Shinomiya T. 1993. Regulation of pyocin genes in Pseudomonas aeruginosa by positive (prtN) and negative (prtR) regulatory genes. J Bacteriol 175:1257–63.

14. Penterman J, Singh PK, Walker GC. 2014. Biological cost of pyocin production during the SOS response in Pseudomonas aeruginosa. J Bacteriol 196:3351–9.

15. Sun Z, Shi J, Liu C, Jin Y, Li K, Chen R, Jin S, Wu W. 2014. PrtR homeostasis contributes to Pseudomonas aeruginosa pathogenesis and resistance against Ciprofloxacin. Infect Immun.

16. Behrens HM, Lowe ED, Gault J, Housden NG, Kaminska R, Weber TM, Thompson CMA, Mislin GLA, Schalk IJ, Walker D, Robinson C V., Kleanthous C. 2020. Pyocin S5 Import into Pseudomonas aeruginosa Reveals a Generic Mode of Bacteriocin Transport. MBio 11:1–16.

17. Denayer S, Matthijs S, Cornelis P. 2007. Pyocin S2 (Sa) kills Pseudomonas aeruginosa strains via the FpvA type I ferripyoverdine receptor. J Bacteriol 189:7663–7668.

18. McCaughey LC, Josts I, Grinter R, White P, Byron O, Tucker NP, Matthews JM, Kleanthous C, Whitchurch CB, Walker D. 2016. Discovery, characterization and in vivo activity of pyocin SD2, a protein antibiotic from Pseudomonas aeruginosa. Biochem J 473:2345–2358.

19. Sano Y, Kobayashi M, Kageyama M. 1993. Functional domains of S-type pyocins deduced from chimeric molecules. J Bacteriol 175:6179–6185.

20. White P, Joshi A, Rassam P, Housden NG, Kaminska R, Goult JD, Redfield C, McCaughey LC, Walker D, Mohammed S, Kleanthous C. 2017. Exploitation of an iron transporter for bacterial protein antibiotic import. Proc Natl Acad Sci U S A 114:12051–12056.

21. Chauleau M, Mora L, Serba J, de Zamaroczy M. 2011. FtsH-dependent processing of RNase colicins D and E3 means that only the cytotoxic domains are imported into the cytoplasm. J Biol Chem 286:29397–407.

22. Kleanthous C. 2010. Swimming against the tide: progress and challenges in our understanding of colicin translocation. Nat Rev Microbiol 8:843–8.

23. Mora L, de Zamaroczy M. 2014. In vivo processing of DNase colicins E2 and E7 is required for their import into the cytoplasm of target cells. PLoS One 9:e96549.

24. Parret AHA, De Mot R. 2002. Bacteria killing their own kind: novel bacteriocins of Pseudomonas and other gamma-proteobacteria. Trends Microbiol 10:107–12.

25. Kühlmann UC, Moore GR, James R, Kleanthous C, Hemmings AM. 1999. Structural parsimony in endonuclease active sites: Should the number of homing endonuclease families be redefined? FEBS Lett 463:1–2.

26. Walker DC, Georgiou T, Pommer AJ, Walker D, Moore GR, Kleanthous C, James R. 2002. Mutagenic scan of the H-N-H motif of colicin E9: implications for the mechanistic enzymology of colicins, homing enzymes and apoptotic endonucleases. Nucleic Acids Res 30:3225–34.

27. Cascales E, Buchanan SK, Duché D, Kleanthous C, Lloubès R, Postle K, Riley M, Slatin S, Cavard D. 2007. Colicin biology. Microbiol Mol Biol Rev 71:158–229.

28. Doudeva LG, Huang H, Hsia K, Shi Z, Li C, Shen Y, Cheng Y, Yuan HS. 2006. Crystal structural analysis and metal-dependent stability and activity studies of the ColE7 endonuclease domain in complex with DNA/Zn^2+^ or inhibitor/Ni^2+^. Protein Sci 15:269–80.

29. Pommer AJ, Cal S, Keeble AH, Walker D, Evans SJ, Kühlmann UC, Cooper A, Connolly BA, Hemmings AM, Moore GR, James R, Kleanthous C. 2001. Mechanism and cleavage specificity of the H-N-H endonuclease colicin E9. J Mol Biol 314:735–49.

30. Kleanthous C, Kühlmann UC, Pommer AJ, Ferguson N, Radford SE, Moore GR, James R, Hemmings AM. 1999. Structural and mechanistic basis of immunity toward endonuclease colicins. Nat Struct Biol 6:243–252.

31. Ko TP, Liao CC, Ku WY, Chak KF, Yuan HS. 1999. The crystal structure of the DNase domain of colicin E7 in complex with its inhibitor Im7 protein. Structure 7:91–102.

32. Maté MJ, Kleanthous C. 2004. Structure-based analysis of the metal-dependent mechanism of H-N-H endonucleases. J Biol Chem 279:34763–9.

33. Turano H, Gomes F, Barros-Carvalho GA, Lopes R, Cerdeira L, Netto LES, Gales AC, Lincopan N. 2017. Tn6350, a novel transposon carrying pyocin S8 genes encoding a bacteriocin with activity against carbapenemase-producing Pseudomonas aeruginosa. Antimicrob Agents Chemother 61:1–4.

34. Sano Y, Kageyama M. 1993. A novel transposon-like structure carries the genes for pyocin AP41, a Pseudomonas aeruginosa bacteriocin with a DNase domain homology to E2 group colicins. Mol Gen Genet 237:161–70.

35. Wallis R, Reilly A, Barnes K, Abell C, Campbell DG, Moore GR, James R, Kleanthous C. 1994. Tandem overproduction and characterisation of the nuclease domain of colicin E9 and its cognate inhibitor protein Im9. Eur J Biochem 220:447–54.

36. Pommer AJ, Kühlmann UC, Cooper A, Hemmings AM, Moore GR, James R, Kleanthous C. 1999. Homing in on the role of transition metals in the HNH motif of colicin endonucleases. J Biol Chem 274:27153–60.

37. Hsia K-C, Chak K-F, Liang P-H, Cheng Y-S, Ku W-Y, Yuan HS. 2004. DNA binding and degradation by the HNH protein ColE7. Structure 12:205–14.

38. Huang H, Yuan HS. 2007. The conserved asparagine in the HNH motif serves an important structural role in metal finger endonucleases. J Mol Biol 368:812–21.

39. Kühlmann UC, Pommer AJ, Moore GR, James R, Kleanthous C. 2000. Specificity in protein-protein interactions: the structural basis for dual recognition in endonuclease colicin-immunity protein complexes. J Mol Biol 301:1163–78.

40. Ku W-Y, Liu Y-W, Hsu Y-C, Liao C-C, Liang P-H, Yuan HS, Chak K-F. 2002. The zinc ion in the HNH motif of the endonuclease domain of colicin E7 is not required for DNA binding but is essential for DNA hydrolysis. Nucleic Acids Res 30:1670–8.

41. Pommer AJ, Wallis R, Moore GR, James R, Kleanthous C. 1998. Enzymological characterization of the nuclease domain from the bacterial toxin colicin E9 from Escherichia coli. Biochem J 334 (Pt 2:387–92.

42. Chassaing B, Cascales E. 2018. Antibacterial Weapons: Targeted Destruction in the Microbiota. Trends Microbiol 26:329–338.

43. Llamas MA, Imperi F, Visca P, Lamont IL. 2014. Cell-surface signaling in Pseudomonas: stress responses, iron transport, and pathogenicity. FEMS Microbiol Rev 38:569–97.

44. Oluyombo O, Penfold CN, Diggle SP. 2019. Competition in biofilms between cystic fibrosis isolates of pseudomonas aeruginosa is shaped by R-pyocins. MBio 10:1–13.

45. Drozdetskiy A, Cole C, Procter J, Barton GJ. 2015. JPred4: a protein secondary structure prediction server. Nucleic Acids Res 43:W389–94.

46. Kabsch W. 2010. XDS. Acta Crystallogr Sect D Biol Crystallogr 66:125–132.

47. Winn MD, Ballard CC, Cowtan KD, Dodson EJ, Emsley P, Evans PR, Keegan RM, Krissinel EB, Leslie AGW, McCoy A, McNicholas SJ, Murshudov GN, Pannu NS, Potterton EA, Powell HR, Read RJ, Vagin A, Wilson KS. 2011. Overview of the *CCP*4 suite and current developments. Acta Crystallogr Sect D Biol Crystallogr 67:235–242.

48. McCoy AJ, Grosse-Kunstleve RW, Adams PD, Winn MD, Storoni LC, Read RJ. 2007. Phaser crystallographic software. J Appl Crystallogr 40:658–674.

49. Emsley P, Cowtan K. 2004. Coot: Model-building tools for molecular graphics. Acta Crystallogr Sect D Biol Crystallogr 60:2126–2132.

50. Vagin AA, Steiner RA, Lebedev AA, Potterton L, McNicholas S, Long F, Murshudov GN. 2004. REFMAC5 dictionary: Organization of prior chemical knowledge and guidelines for its use. Acta Crystallogr Sect D Biol Crystallogr 60:2184–2195.

